# Value-based Decision Making Takes Place in the Action Domain in the Prefrontal Cortex

**DOI:** 10.1101/444646

**Authors:** Zhongqiao Lin, Chechang Nie, Yuanfeng Zhang, Yang Chen, Tianming Yang

## Abstract

Value-based decision making is a process in which humans or animals maximize their gain by selecting appropriate options and performing the corresponding actions to acquire them. Whether the evaluation process of the options in the brain can be independent from their action contingency has been hotly debated. To address the question, we trained rhesus monkeys to make decisions by integrating evidence and studied whether the integration occurred in the stimulus or the action domain in the brain. After the monkeys learned the task, we recorded both from the orbitofrontal (OFC) and dorsolateral prefrontal (DLPFC) cortices. We found that the OFC neurons encoded the value associated with the single piece of evidence in the stimulus domain. Importantly, the representations of the value in the OFC was transient and the information was not integrated across time for decisions. The integration of evidence was observed only in the DLPFC and only in the action domain. We further used a neural network model to show how the stimulus-to-action transition of value information may be computed in the DLPFC. Our results indicated that the decision making in the brain is computed in the action domain without an intermediate stimulus-based decision stage.

## Introduction

We often have to make choices between different options based on their value. Naturally, the choices are tied to actions that are used to acquire the options. In many cases, the evaluation of each option is a complex process in which one has to take consideration of multiple pieces of information and integrate them. Although our subject experience may suggest that we only carry out an action to substantiate a decision after it is made, many studies have shown that the neurons at different levels of motor pathway may reflect the decision-making process long before (Basso and Wurtz, 1998; Cisek and Kalaska, 2005; Gold and Shadlen, 2000; Hernández et al., 2002; Kim and Shadlen, 1999; Romo et al., 2004; Shadlen and Newsome, 2001). These studies inspired the hypothesis that decision making is implemented in the brain as an action selection process in which values for competing actions are calculated and compared. This would allow actions to be carried out as soon as the decisions are made, which makes sense as animals in the real world need to make responses quickly to survive.

It has, however, been argued, that some decisions are made in the brain with an intermediate stage where the decision-making and value computation process is completely dissociated from their motor contingency. Recent investigations seem to suggest that during economic decisions, there is a representation of value in the brain that is independent from the motor contingency (Cai and Padoa-Schioppa, 2014; Chen and Stuphorn, 2015; Padoa-Schioppa and Assad, 2006; Wallis and Miller, 2003). For example, Padoa-Schioppa and colleagues reported that a group of the orbitofrontal (OFC) neurons encoded the value of the chosen option regardless of the direction of the eye movement used by the animals to indicate their choice (Padoa-Schioppa and Assad, 2006, 2007; Padoa-Schioppa and Conen, 2017). Based on these studies, it was proposed that the brain calculates the values of competing options during decision making independent from their associated actions and the OFC is a candidate brain area for computing value in an action-independent manner (Padoa-Schioppa, 2007, 2011). In addition, lesions in the OFC were shown to lead to deficits in stimulus-value updating but not action-value updating (Rudebeck et al., 2008).

These experiments that examined the value representation in the OFC were based on the behavior tasks with a distinct aspect. In these tasks, the decisions were based on rather simple stimulus-reward associations (Cai and Padoa-Schioppa, 2014; Kennerley et al., 2011; Padoa-Schioppa and Assad, 2006; Raghuraman and Padoa-Schioppa, 2014), which could be computed quickly and left little room for motor preparation. If decisions have to be calculated in an extended process as the ones investigated in many perceptual decision making studies (Gold and Shadlen, 2000; Hernández et al., 2002; Kim and Shadlen, 1999; Shadlen and Newsome, 2001), it is unclear whether they could be carried out entirely independent of their action contingencies.

Here, we investigate the question with a task in which monkeys had to make choices between two colored targets associated with probabilistic rewards. The reward probability of each color was indicated by a sequence of simple shape pictures that served as visual cues. The monkeys had to combine information from these shapes to calculate the reward probabilities and figure out the more rewarding target. Their choices were indicated with saccadic eye movements. Critically, both red and green targets could appear on either the left or the right, randomly chosen by the computer in each trial. The evidence was provided regarding to the target color, independent from eye movement directions. Because the saccade circuitry in the brain uses spatial coordinates, we investigate how the value information regarding to the color (stimulus-based) is transformed into the value information regarding to the spatial location (action-based) in the brain to form decisions. We recorded single unit activities from the DLPFC and the OFC. We found the OFC neurons encoded evidence associated with each single piece of evidence, but only in the stimulus domain. The integration of evidence was only represented in the DLPFC and only in the action domain. These results argue against the role of the OFC in value computation during decision making and suggest that the decision for actions is calculated in the brain without an intermediate stimulus-based decision stage.

## Results

The experiments and analyses were done with two macaque monkeys. We present the results here based on the data combined from both monkeys; the individual monkeys’ results are consistent and may be found in the supplementary figures.

### Behavior

We trained the monkeys to perform a probabilistic reasoning task (Figure 1A). The task was slightly modified from a previous study (Yang and Shadlen, 2007). In each trial, the animals were shown a sequence of four shapes, drawn randomly with replacement from a pool of ten shapes. Each shape was assigned with a weight. The sum of the weights was the log odds between the reward probabilities of the red target and the green target (Eqs. 1 and 2). Thus, the shapes with positive weights indicated that the red target had the larger reward probability, whereas the shapes with negative weights indicated that the green target had the larger reward probability. The monkeys reported their choice with a saccadic eye movement toward the chosen target. The reward was delivered probabilistically based on the summed weight of the shapes in the sequence. The red and the green targets were randomly placed either on the left or the right side of the screen.

**Figure 1.**
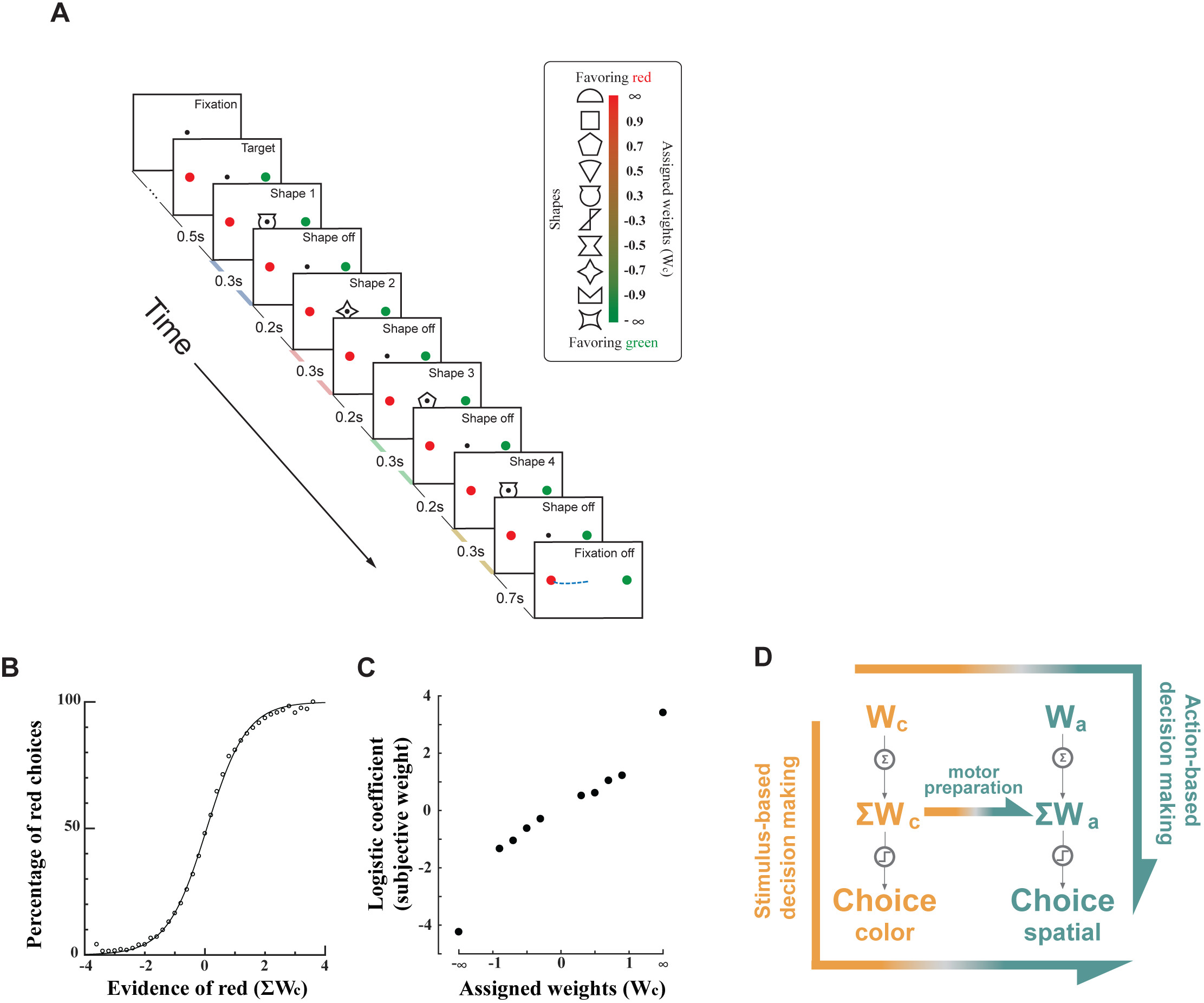
**(A)** Task design. Four shapes were presented sequentially on the computer screen while the monkey fixated at the FP. After the FP was turned off, the monkey made a saccade to either the red or green choice target. The shapes were selected randomly with replacement in each trial from a set of ten. Each of them was assigned with a unique weight (inset). The reward probability was calculated by the sum of the weights associated with the four shapes. Positive weights indicated the red target had a reward probability larger than 0.5. The blue, red, green, and yellow shadings along the time axis indicate the stimulus representation periods 1~4. **(B)** Monkey performance. The percentage of red choices is plotted against the summed weight of the four shapes (black dots). The curve is a fitted logistic function. **(C)** Subjective weights. We used a logistic regression to assess the effects of each shape on the monkeys’ choice and define the coefficients as the subjective weights. Positive weights indicate a tendency to choose the red target. **(D)** Two competing decision-making hypotheses. In stimulus-based decision making, the information regarding to color (*Wc*) is first integrated to the Σ*Wc*, which is then used to generate the action-independent color choice and finally translated into the actions. In contrast, the action-based decision making hypothesis assumes that the *Wc* is first transformed into the action domain (*Wa*), which is then accumulated into the Σ*Wa* and forms the choice and actions. In addition, the stimulus-based decision making may be complemented by a motor preparation process, in which the Σ*Wa* is calculated from the Σ*Wc* during the decision making process.

Both monkeys learned to perform the task (Figure 1B and Supplementary Figure 1A,B). The monkeys chose the green target more often when the summed color weight was negative, and the red target more often as the summed weight was positive. To assess how each shape affected the monkeys’ choices, we applied a logistic regression with the number of appearances of each shape as the regressors (Eq. 4). The regression coefficients showed the same rank order as the assigned weights, indicating the monkey assigned appropriate weights to the shapes (Figure 1C and Supplementary Figure 1C,D).

The task design allowed us to distinguish the stimulus-based and action-based decision-making processes in the brain. The shape weights indicated the reward probability of each target color. Both colors could appear on the left or the right. Thus, the value associated with the color is orthogonal to the value associated to the eye movement direction. Because the eye movement circuitry in the brain carries out the motor commands using spatial coordinates, the outcome of the decision making has to be transformed into the spatial domain eventually. The critical question is whether the decision process of where to move the eyes, i.e. the integration process of the evidence, is carried out in the action or the stimulus domain. If it is the former, we should observe that the evidence associated with each shape is first transformed into the action domain and then integrated. Thus, the representation of the integrated evidence should only be found in the action domain in the brain. If it is the latter, we would observe the representation of the integrated evidence in the stimulus (i.e. color) domain, which would be transformed into the action domain at a later stage. An alternative scenario of the stimulus-based decision making is that there is a separate motor preparation process during the stimulus-based decision making, in which the action value is calculated from the stimulus value. In this scenario, the representation of the integrated evidence in both the stimulus and the action domains may be observed, but not necessarily the representation of the individual piece of evidence in the action domain. Therefore, with this behavior paradigm, we may find out how the decision making unfolds in the brain by investigating in which domain neurons in the prefrontal circuitry encode evidence (Figure 1C).

For the convenience of discussion of the neuron data below, in any given epoch, we use *Wc* to refer to the weight associated with the particular shape appearing in that epoch in the color domain, *Wa* the weight associated with the single shape in the action domain, ∑*Wc* the sum of the weights of the shapes that have appeared so far in the color domain, and ∑*Wa* the sum of the weights of the shapes that have appeared so far in the action domain.

### Example Neurons

We recorded single unit activities from the OFC and the DLPFC. **Figure 2** shows two representative example units from each area, respectively. The example neuron from the DLPFC showed a response pattern similar to what was previously described in the LIP (Kira et al., 2015; Yang and Shadlen, 2007). The neuron’s responses ramped up or down as the evidence grew in favor or against the target associated with its preferred eye movement direction. When we sorted the trials into quartiles according to the Σ*Wa* in each epoch, we observed greater responses when the Σ*Wa* was larger (Figure 2A). In contrast, the example neuron from the OFC did not have a clear ramping activity pattern. Its responses were instead modulated by the *Wc*. When we sorted the trials by the weight of the shape presented in each epoch regarding to color, we found the neuron’s responses were greater when the evidence was more in favor of the red target (Figure 2B).

**Figure 2.**
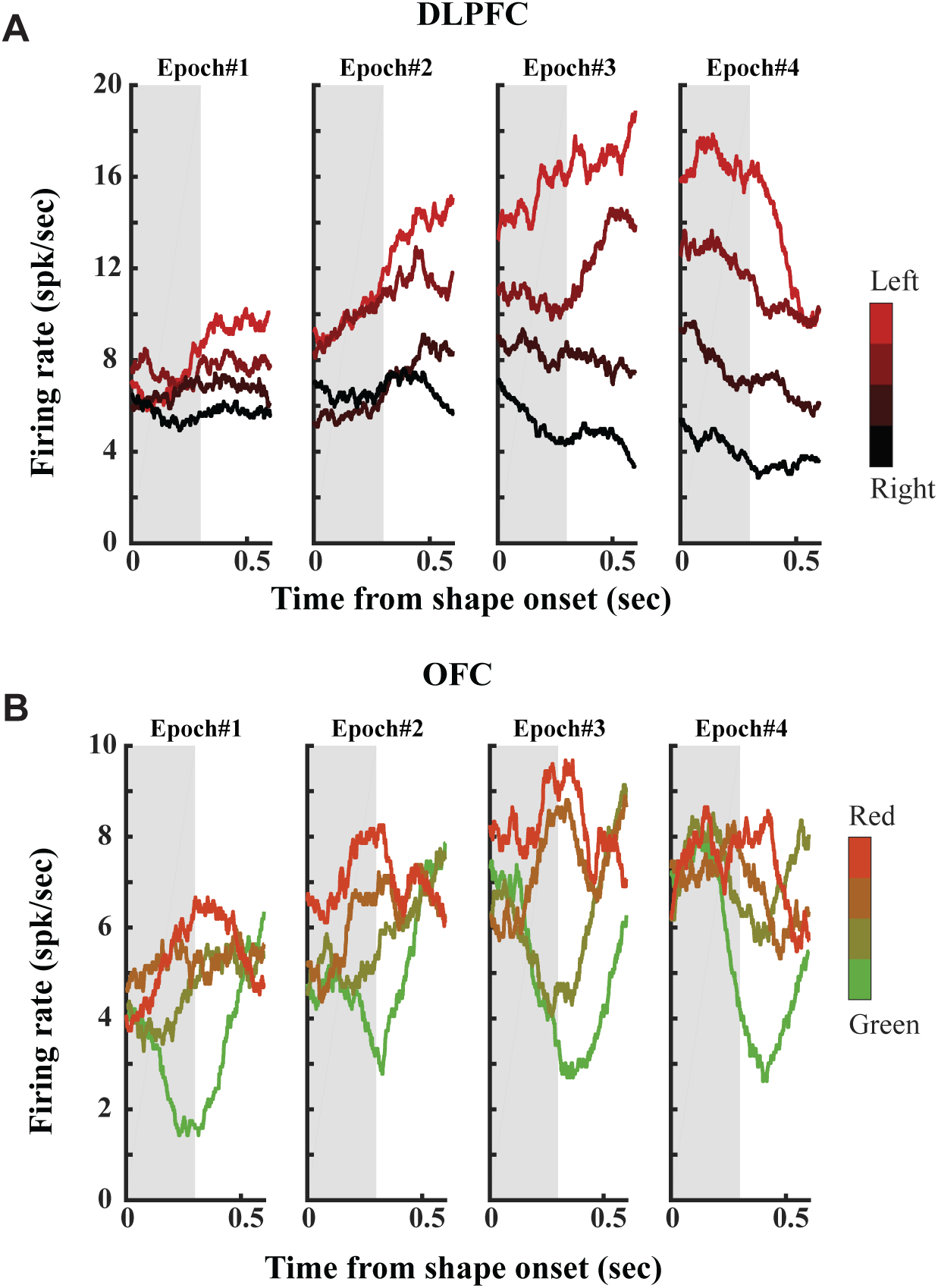
Example neurons. **(A)** The activity of a DLPFC neuron encoded the Σ*Wa*. Response averages are aligned to the shape onsets and extended 100 ms into the next shape period. The trials are divided into quartiles by the Σ*Wa* (indicated by the redness) and the neuron’s response averages are computed. The width indicates the s.e.m. across trials. The gray shades indicate the period when the shape was displayed (300 ms). **(B)** The activity of an OFC neuron encoded the *Wc*. Response averages are aligned to the shape onsets and extended 100 ms into the next shape period. The trials are divided into quartiles by the *Wc* (indicated by the color) and the neuron’s response averages are computed. The width indicates the s.e.m across trials. The gray shades indicate the shape presentation period (300ms).

The example neurons showed that the neurons in both the OFC and the DLPFC encoded relevant information for decision making. To fully appreciate the roles that the two areas play in decision making, we recorded activities of 277 cells from the OFC (121 and 156 from monkeys K and E, respectively) and of 384 cells from the DLPFC (170 and 214 from monkeys K and E, respectively).

### Population Analyses: Choices

First, we asked the question whether the choice outcome was encoded in the two brain areas. This would provide us clues whether they were involved in decision making. Just as the weights are, the choice outcome could also be in the color domain and in the action domain. We looked at them separately.

We sorted all trials according to each neuron’s preference of either color or eye movement direction and compared the neurons’ responses between the choice outcomes (Figure 3). We found that the OFC neurons barely signaled the monkeys choice in either the color or the action domain during the stimulus presentation period (Figure 3A). The DLPFC neurons, however, were strongly modulated by the monkeys’ choice outcome regarding to the eye movement direction (Figure 3B). The modulation became significant at 280 ms after the 3^rd^ shape epoch and was maintained till the end of the trial. The DLPFC neurons also showed a difference between their responses to the two color choices, although in a much more modest and less consistent manner. The representation of the color choice in the DLPFC did not precede that of the spatial choice.

**Figure 3.**
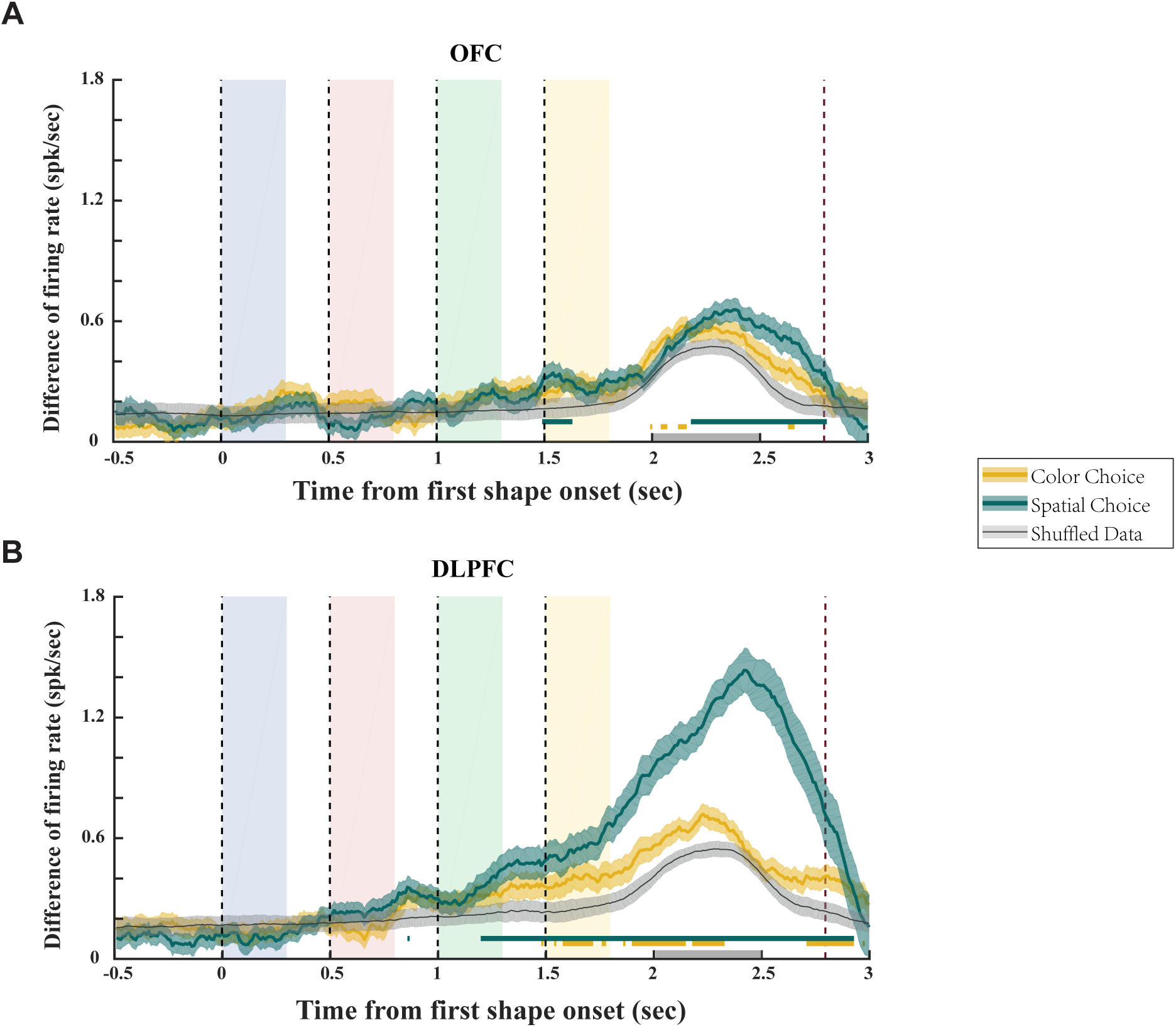
Representation of choice. **(A)** The average firing rate difference of the OFC neurons in trials with different choice outcomes. Green: spatial choice; yellow: color choice; black: shuffled data. The green and yellow shaded areas indicate the s.e.m., and the grey shaded area indicates the s.d. of the 100 shuffles. The green and yellow horizontal lines mark the periods in which the spatial or color choice curve is significantly different from the shuffled data (p<0.01, one-way ANOVA). The four color shaded boxes indicate the shape presentation period in the four epochs, with the dashed lines indicating the shape onset. The rightmost dashed line in dark red marks the average saccade time. The grey bar on the horizontal axis represents the period from which the mean firing rates were calculated to define the neurons’ preferred choices. **(B)** The average firing rate difference of the DLPFC neurons for different choices.

### Population Analyses: Stimulus Weights

We further studied whether the neuronal responses in the two areas captured different stages of decision making. To understand the complete picture of how neurons in the OFC and the DLPFC contributed to the task, we resorted to the regression analyses to find out how each relevant variable in the task may explain the populational responses in each area. More specifically, we performed linear regressions of each neuron’s responses on the *Wc*, *Wa*, Σ*Wc*, and Σ*Wa* from each of the four epochs in a trial and looked at how well the neurons’ responses were explained by each variable from each epoch. We used the shuffled data to test for significance (See Methods). This approach allows us to consider neurons with different tuning properties together and estimate how information is encoded by each population.

Among the four variables, OFC prominently encoded the *Wc* (Figure 4A). The encoding, however, was not sustained. It reached significance level on average 192 ms across the 4 epochs after the shapes’ onset, and disappeared shortly after the next shape appeared. There was little overlapping between two consecutive shapes, suggesting a lack of information integration. Although the regression showed an apparent representation of the Σ*Wc* by the OFC (Figure 4C), it was weak and also transient. The fact that the OFC did not encode the color choice outcome further suggested this apparent representation of the Σ*Wc* was a statistical artifact, which might be due to the inherent correlation between the *Wc* and the Σ*Wc*. Importantly, the encoding of the weights was restricted to the stimulus domain and not observed in the action domain (Figure 4B,D). The individual monkey analyses also yielded consistent results (Supplementary Figure S2).

**Figure 4.**
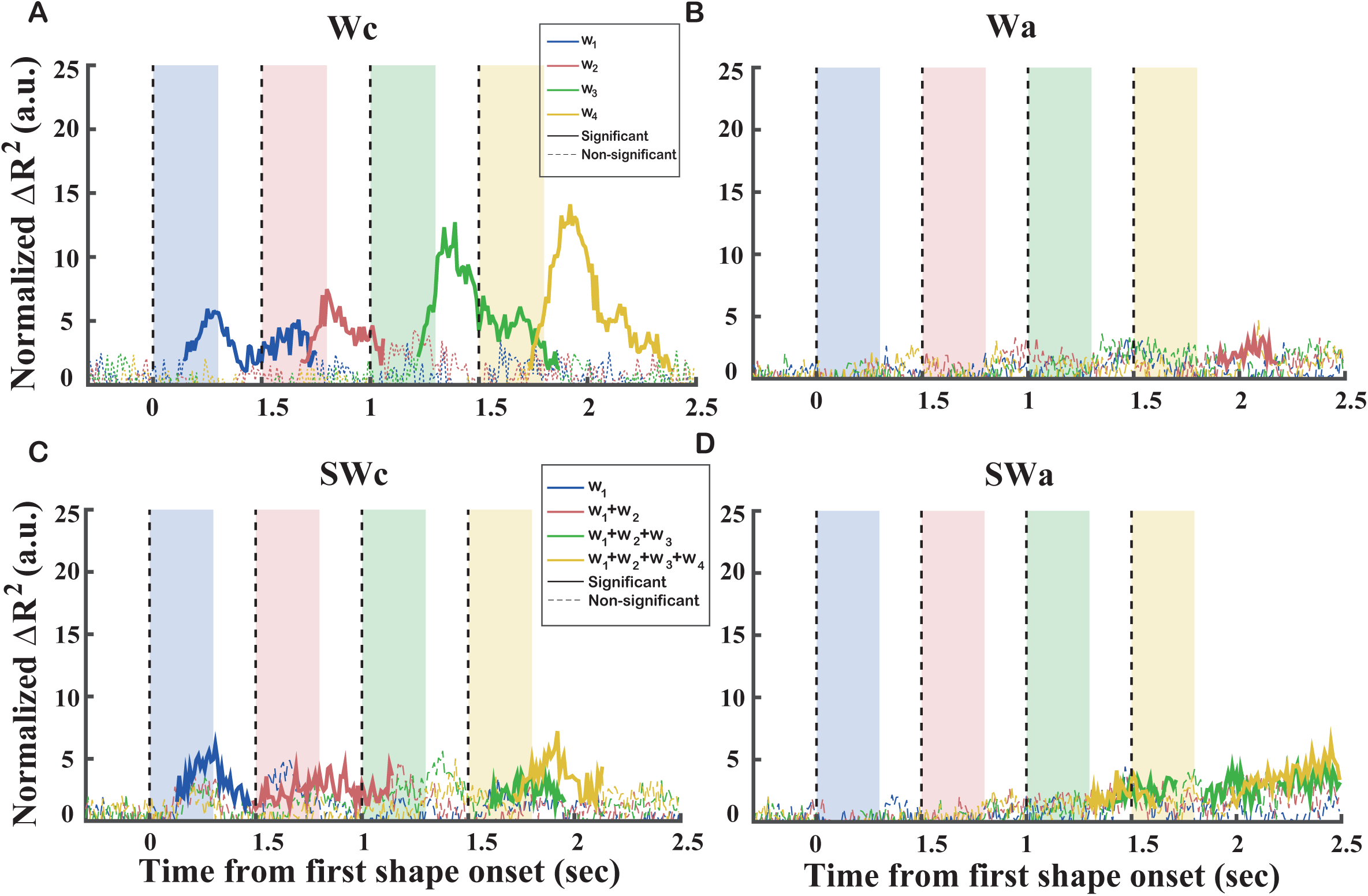
Representations of the shape weights in the OFC. **(A-D)** The normalized explained variance of *Wc*, *Wa*, Σ*Wc*, and Σ*Wa* of the OFC neurons. Blue, red, green, and yellow indicate the 1^st^, 2^nd^, 3^rd^, and 4^th^ epoch, respectively. The solid sections of each curve indicate significance (p<0.05 with multiple comparison corrections), and the dashed sections are not significant.

In contrast to the OFC neurons, DLPFC neurons exhibited very different response patterns (Figure 5). First of all, both variables in the action domain (*Wa* and Σ*Wa*) were strongly represented in the DLPFC (Figure 5B,D). In addition, their representations were sustained till the end of the trial. The representation of the *Wa* exhibited a relatively flat pattern, while the representation of the Σ*Wa* showed clear ramping. Such a pattern is a signature of the integration of information across different epochs. The DLPFC neurons were also found to encode the *Wc*, although the encoding was much weaker than that of the *Wa* (Figure 5A). In addition, the encoding of the *Wc* appeared to be transient in a similar fashion as in the case of the OFC, suggesting a lack of integration in the stimulus domain. Consistent with this observation, the encoding of the Σ*Wc* in the DLPFC was very weak, if it existed at all (Figure 5C). These results were further supported by the individual monkey analyses (Supplementary Figure S3).

**Figure 5.**
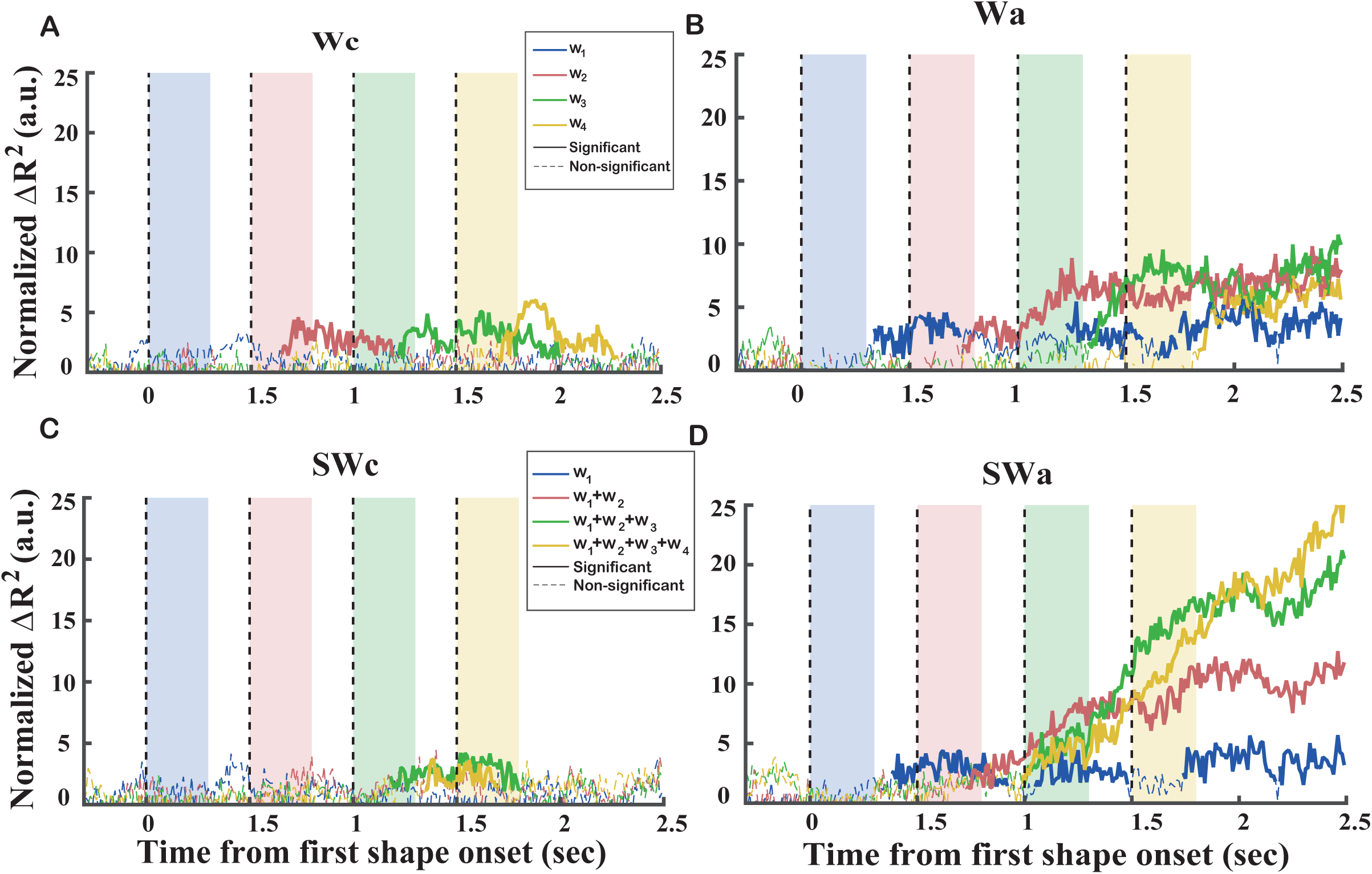
Representations of the shape weights in the DLPFC. **(A-D)** The normalized explained variance of *Wc*, *Wa*, Σ*Wc*, and Σ*Wa* of the DLPFC neurons. Blue, red, green, and yellow indicate the 1^st^, 2^nd^, 3^rd^, and 4^th^ epoch, respectively. The solid sections of each curve indicate significance (p<0.05 with multiple comparison corrections), and the dashed sections are not significant.

### LASSO

The regression analyses carried out above were done separately for each variable and for each epoch (See Methods). This might cause concerns, because the weight terms and the summed weight terms were linearly dependent. Therefore, the positive findings above should be interpreted cautiously. To alleviate this problem and confirm our findings, we applied LASSO on a model containing all four weight variables in the previous analyses, as well as the two choice variables: one for the color and one for the eye movement direction. Because of the penalty term in the LASSO, it tends to use a smaller number of variables to explain the data (Tibshirani, 1996). Therefore, if, for example, a neuron’s responses could be well explained by Σ*Wa*, the LASSO analysis would assign the *Wa* terms with only small weights. Thereby, the LASSO analysis may provide us further information on whether a particular neuron population encoded the weights or the summed weights.

To compare the results from different neuron populations and across different variables, we calculated the normalized absolute standard regression coefficients (|*SRβ*|) to evaluate the effects of each individual variable (see **Methods**), which is analogous to the normalized Δ*R*^2^ used in linear regression analyses. They are plotted in **Figure 6** for the OFC and in **Figure 7** for the DLPFC, and in Supplementary Figure S4 and S5 for individual monkeys. Note that we omitted the trace of the first epoch in all the plots of the summed weights, because they were the same as that of the single weights by design (see **Methods**).

**Figure 6.**
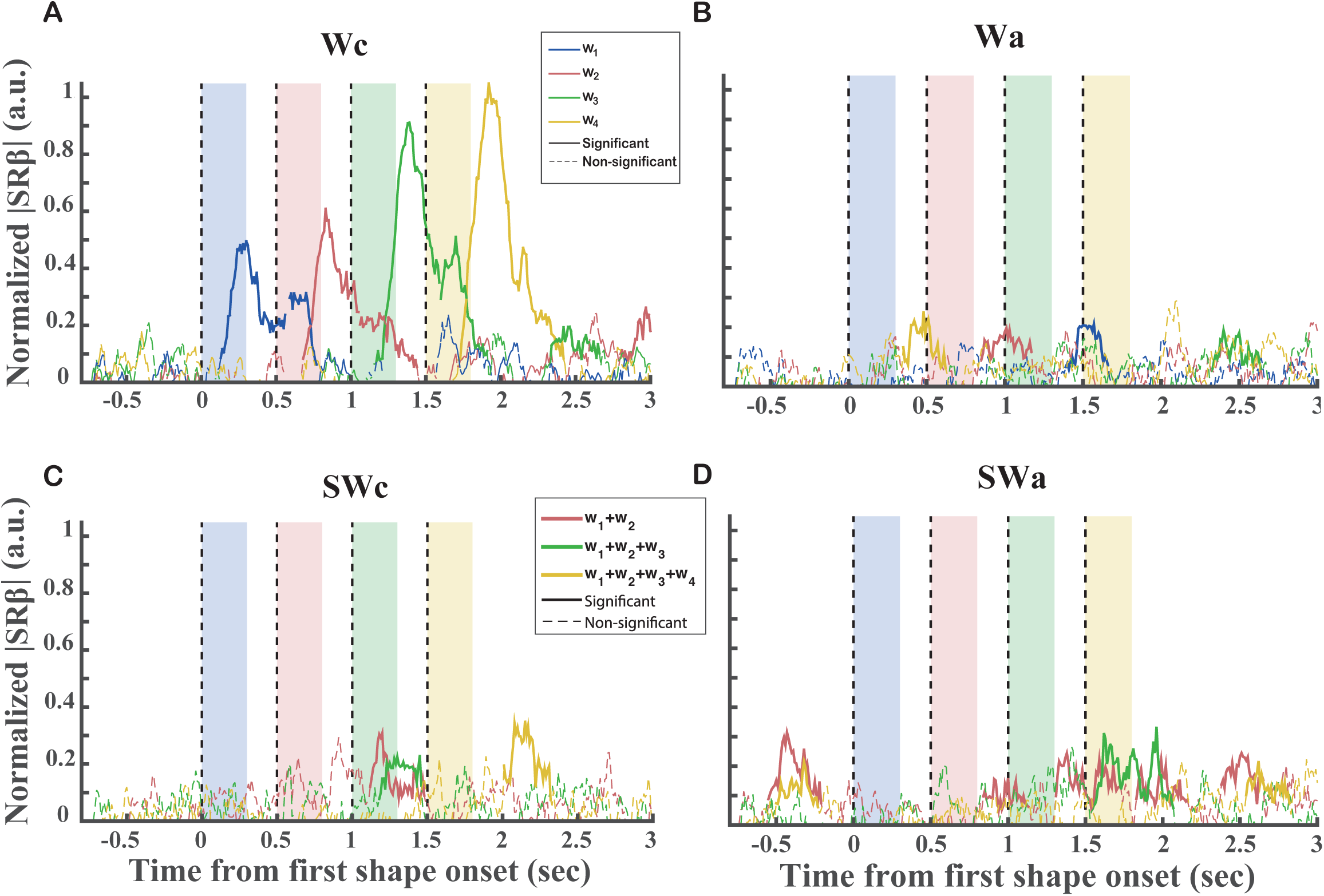
Representations of the shape weights in the OFC with LASSO. **(A-D)** The normalized |SRβ| of *Wc*, *Wa*, Σ*Wc*, and Σ*Wa* of the OFC neurons. Blue, red, green, and yellow indicate the 1^st^, 2^nd^, 3^rd^, and 4^th^ epoch, respectively. The solid sections of each curve indicate significance (p<0.05 with multiple comparison corrections), and the dashed sections are not significant. The traces of the Σ*Wc* and Σ*Wa* from the first epoch in were omitted (see **Methods**).

**Figure 7.**
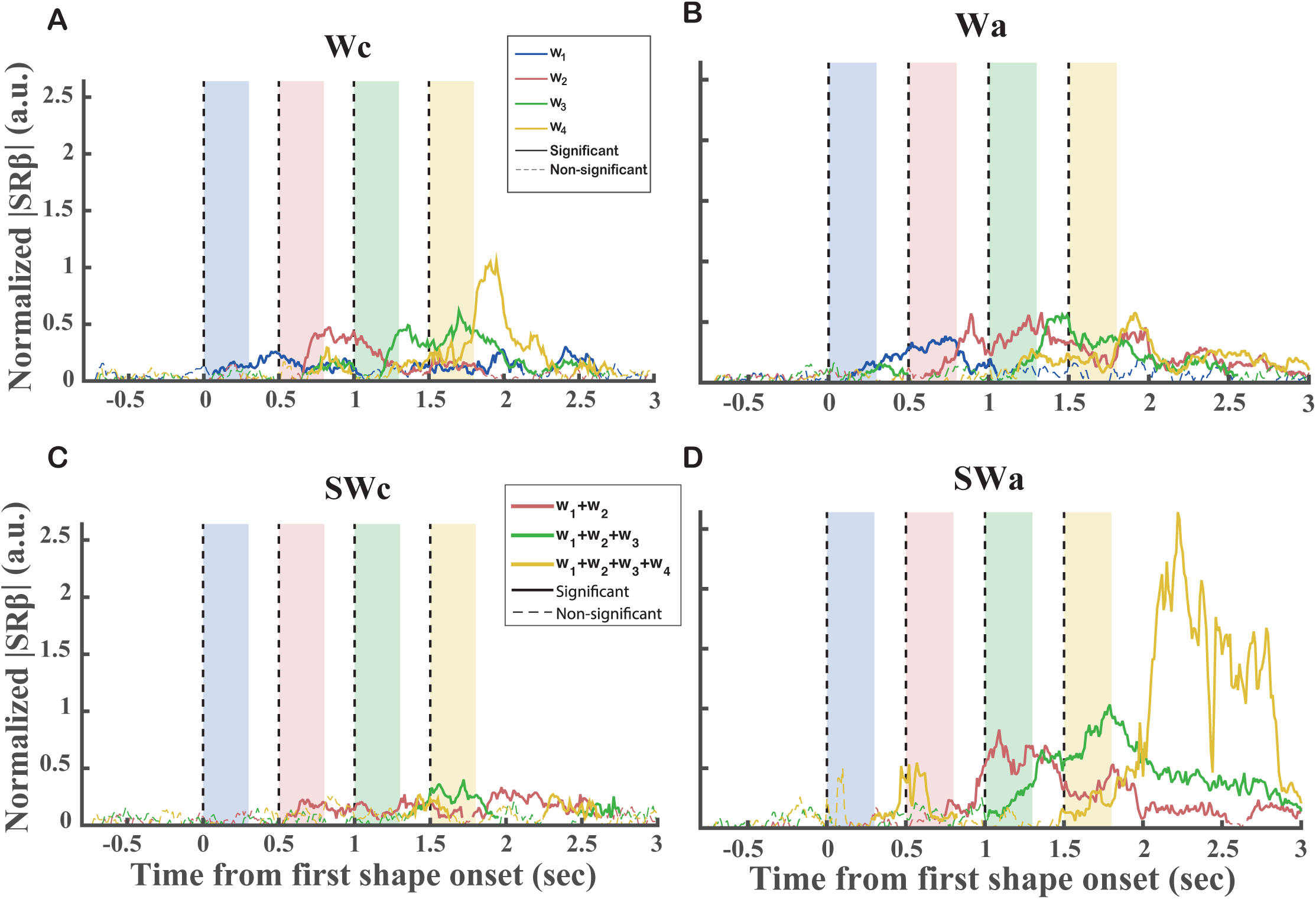
Representations of the shape weights in the DLPFC with LASSO. **(A-D)** The normalized |SRβ| of *Wc*, *Wa*, Σ*Wc*, and Σ*Wa* of the OFC neurons. Blue, red, green, and yellow indicate the 1^st^, 2^nd^, 3^rd^, and 4^th^ epoch, respectively. The solid sections of each curve indicate significance (p<0.05 with multiple comparison corrections), and the dashed sections are not significant. The traces of the Σ*Wc* and Σ*Wa* from the first epoch in were omitted (see **Methods**).

The results further confirmed our findings with the linear regressions. Consistent with our analyses above, the OFC population were tuned for the *Wc* only (Figure 6A). The LASSO also revealed very little representation of the Σ*Wc* in the OFC, which agrees with our speculations on the possible artifacts in the regression analyses. Also consistent with the results of simple linear regression analysis, the DLPFC population encoded the *Wa*, Σ*Wa*, and *Wc* (Figure 7A, B, D). The peaks of tuning that reached significance in these plots appeared sequentially in the proper order. On the contrary, the normalized |*SRβ*|s of the Σ*Wc* were not only more noisy and weaker, but were also often misplaced (Figure 7C). One may notice interesting differences between the results from the LASSO and the simple linear regressions, which we will discuss later in the Discussion.

### Stimulus-to-Action Transition

The recording experiments suggested that the transition between the stimulus-based signal and the action-based signal occurred at the single-weight stage in the DLPFC. To understand how this transition may be implemented by a neural network, we created a simple neural network model that contained a hidden layer in which units receive inputs of the *Wc* and the spatial configuration (Figure 8A). In the model, the spatial configuration refers to whether the red target is on the left or the green target is on the left and takes the values of ±1. If we define the *Wa* as positive when the eye movement is toward the left, the *Wa* should equal to the spatial configuration times the *Wc*. The network took *Wc* as the input and was trained to transform it into *Wa* as its output.

**Figure 8.**
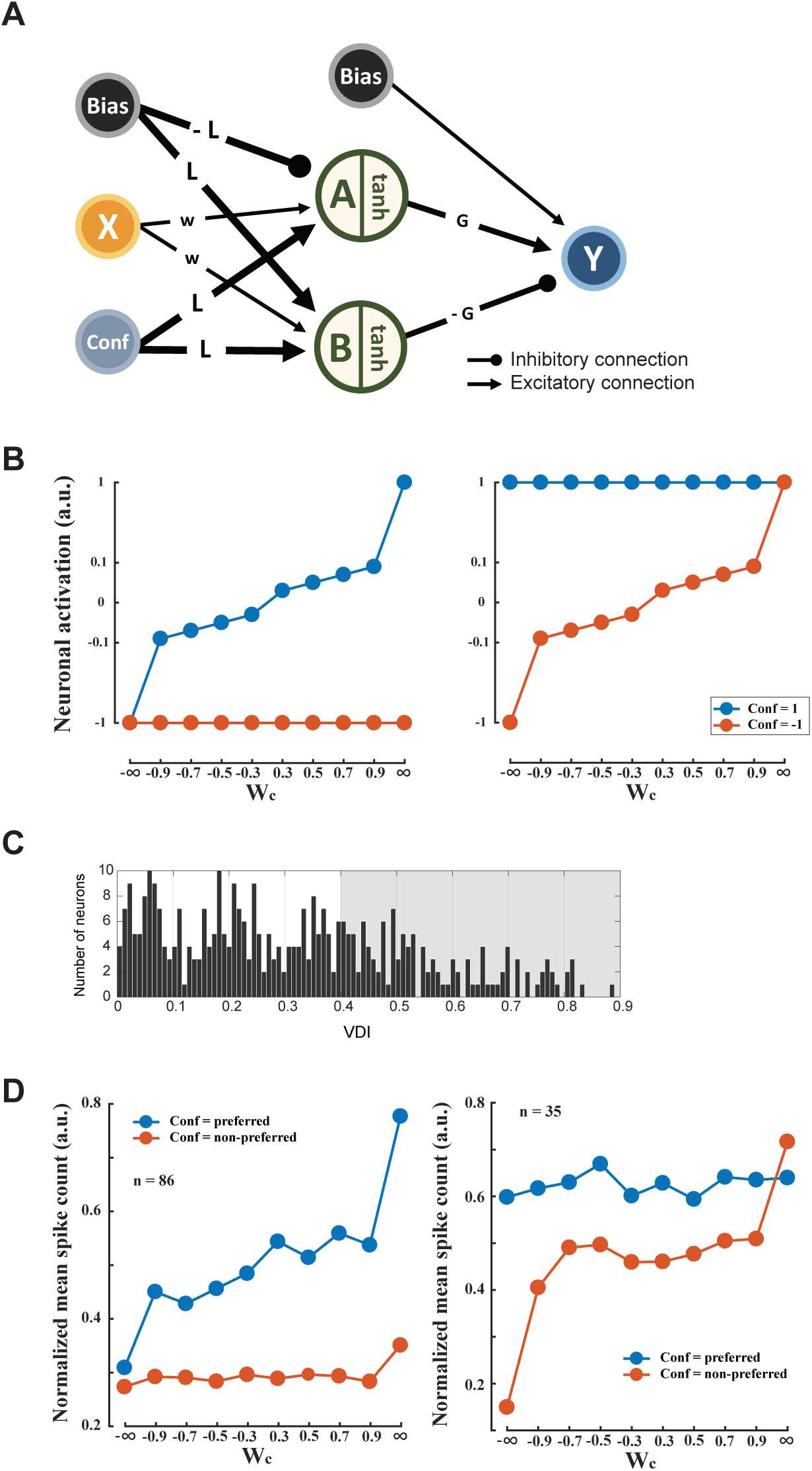
The feedforward network model. Two types of hidden neurons receive inputs from a *Wc* neuron and a spatial configuration neuron. They also receive a bias input. The hidden neurons send projections to the output neuron Y, which encodes *Wa* (see **Methods**). **(B)** The response patterns of the two types of hidden neurons from the model. Each type has an asymmetric response pattern, in which its response under one spatial configuration is flat but not under the other. **(C)** The distribution of VDI of the DLPFC neurons. The shaded area indicates the neurons with VDI greater than 0.4. **(D)**The response patterns of the DLPFC neurons that had the largest variance disparities between the two spatial configurations. The neurons that had the greater responses under the spatial configuration with the larger variance (blue) than that with the smaller variance (red) are plotted in the left panel, and the neurons that had weaker responses under the spatial configuration with the larger variance (red) than that with the smaller variance (blue) are plotted in the right panel.

We found that the hidden layer may be reduced to having only two types of units (See Methods). The activity of the first type of units encoded the *Wc* under only one spatial configuration and was flat across the different shapes under the other configuration. The activity pattern of the other type of units was the opposite. Their activity encoded the *Wc* under the spatial configuration opposite to the one that was encoded by the first type of units (Figure 8B). The output unit pooled the inputs from these two types of units with weights of opposite signs.

It is easy to see that the bias input to one of the hidden units (e.g. A) was cancelled under one configuration (+1), allowing its output to reflect the color weight input.

However, the other hidden unit (B) was saturated under the same condition. The output unit pooled the hidden units’ responses with weights in opposite signs and added a bias term to cancel out the input from the saturated unit. Under the other configuration, the two units were reversed so that now A was saturated and B encoded the weight. The eventual output was the same with this symmetry. The output *Y* ≈ *Conf* * *Wc*, which was just *Wa*.

We reasoned that we would see some DLPFC neurons exhibit similar activity patterns if the DLPFC were where the good-to-action transition occurs. To identify these neurons, we noticed that they should have flat tuning curve in one configuration and a steep tuning curve in the other, leading to a big difference in the response variances under the two spatial configurations. We defined a variance difference index (*VDI*) to quantify this pattern (See Methods). We set a threshold of *VDI* at 0.4, which is equivalent to the situation that the variance in the preferred configuration accounts for 70% of the total variance. Among the total of 384 DLPFC neurons we recorded, 121 neurons were selected (Figure 8C). We further divided them into two groups according to whether they had larger responses under the configuration with more variance or under the configuration with less variance. The mean responses to each shape of the two groups of neurons under the two spatial configurations were plotted in Figure 8D. We found that the two groups of DLPFC neurons showed activity patterns that matched the two neuron types from the network model.

These results suggested that the DLPFC neurons not only encoded single weights in both the stimulus and the action domain, they showed activity patterns that could explain the computation underlying the stimulus-to-action transition.

### Motor Preparation

A variation of the stimulus-based model that may explain the choice signal in the action domain during decision making is that it merely reflects a motor preparation signal based on the integration of information in the stimulus domain. Our results do not support this scenario. For that to be true, the motor preparation signal has to be calculated online based on a decision variable in the stimulus domain, which is the Σ*Wc* in our experiment. However, we did not find evidence that either the OFC or the DLPFC neurons encoded the summed weight in the stimulus domain during decision making. Thus, the integration of information happened only in the action domain in the DLPFC, and it did not arise from an intermediate stimulus-based stage.

## Discussion

Here, we have shown how the stimulus-based and action-based decision-making signals were represented in the OFC and DLPFC. These results supported the hypothesis that the decision for actions is computed in the action domain, and the transition between the stimulus-based and action-based value information occurs in the DLFPC at the stage of single piece of evidence.

### Good-based vs Action-based

Several previous studies argued there are stages in the brain where neural activities reflected an abstract good-based decision independent of the motor outcome (Cai and Padoa-Schioppa, 2014; Chen and Stuphorn, 2015; Padoa-Schioppa and Assad, 2006; Wallis and Miller, 2003). Under scrutiny, their observations were actually compatible with our results. In those studies, the decisions were based on simple stimulus-reward associations. This is analogous to the single color weights assigned to the individual shapes in our study. In this sense, we indeed also observed the representations of good-based weights in both the OFC and the DLPFC, which was interpreted as evidence for good-based decision making in the previous studies. However, in our more sophisticated task design, decisions depended on the integration of information provided by a sequence of stimuli. Because of the large number of possible stimuli combinations, the integration was most likely based on a computation online instead of on established stimulus-reward associations. What we did not observe was this integration of information in the stimulus domain. Based on these results, we believe that the previous interpretations based on simpler task designs need to be updated.

Our results, however, cannot exclude the possibility that there is a separate system in the brain that carries out good-based decision making. Such a system is obviously helpful for the brain to establish stimulus-reward associations. Similar to the dichotomy of the ventral and dorsal pathways in the visual systems, the brain may also have two separate pathways for decision making (Rushworth et al., 2012). Several studies have pointed out that the ventral PFC areas contain a stimulus-based attention system (Bichot et al., 2015; Wardak et al., 2010). Given the fact that attention and decision making are often closely tied, these studies may suggest that the VLPFC may play a similar role in the good-based decision-making circuitry as that of the DLPFC in the action-based. What we would like to conclude based on our results is that decision for action in the brain is not based on an intermediate stage in which the decision based on good is first formed. Instead, it is computed entirely in the action domain.

### Regression and LASSO

One of the challenges of the study is trying to disentangle the encodings of the single weights and the summed weights by the neurons. We achieve this by combining the simple linear regression and LASSO analyses.

The regression analyses with single factors gave the strongest argument when a negative result is found. The negative finds would not be changed when other factors were added into the regression, regardless of their linear dependencies. Thus, the lack of representations of the summed weights in the color domain in the OFC or DLPFC is strongly supported by our regression analyses.

The positive findings from the regression, however, should be taken more cautiously because of the inherent correlation between the single weights and the summed weights. This problem cannot be easily addressed by throwing both in the same regression. Even for a hypothetical neuron that only encodes the summed weights, adding the single weights into the regression can still improve the regression results. This is because the system cannot be perfect. There might be noise in the system, or the monkeys do not calculate the combined weights perfectly linearly. A small overweight or underweight during the integration of information for a particular piece of evidence may be better accounted for by adding single weights into the regression. We used LASSO as a way to balance the contributions between the single weights and the summed weights. The LASSO provided results largely consistent with the simple linear regression analyses and strengthened our conclusions.

The temporal dynamics of the encoding provides additional clues for us to interpret these results. If we believe a neuron combines the information from multiple cues and encode the combined evidence, its encoding for both the single weights and the total weights should be persistent. This is indeed observed in the DLPFC neurons in the action domain. In contrast, the encoding of the single weights in the OFC was only transient and lasted little longer than the presentation of each single cue. The OFC neurons did not maintain the information long enough for integration. Thus, they most likely did not directly calculate the combined evidence.

It is interesting to further compare the subtle differences in the positive findings between the simple linear regression and the LASSO analyses. Note that the representations of *Wa* in the DLPFC described by the LASSO appeared to be transient. This did not mean that the representations of *Wa* were also transient in the DLPFC. Instead, it was due to the fact that the sustained representations of the single weights observed in the linear regressions may be better explained by the summed weights in the LASSO in the later period, suggesting a computing process from the single weights to the summed weights. For similar reasons, the growing choice outcome signal largely cancelled out the ramping-up pattern of the Σ*Wa* observed in the linear regression analysis, so the representation of the Σ*Wa* also appeared to be transient in LASSO. This suggest that the representations of the summed weights in the DLPFC were later replaced by the representation of the binary choice signal. The similarities between the results from the LASSO and the linear regression analyses confirmed our conclusions, while their differences added refined points provided uniquely by each analysis.

### OFC and Value

Many studies have shown that the OFC is important for dynamically updating and tracking stimulus-value association (Morrison et al., 2011; Rolls et al., 1996; Rudebeck et al., 2008, 2017; Thorpe et al., 1983). Here we showed that the OFC did not integrate and update value information during decision making. How shall we reconcile our results with the others?

We believe the key difference here is the stimuli we used in our experiment. In our study, the reward value calculated based on multiple pieces of evidence was not directly associated with a concrete stimulus. We used a set of 10 shapes that comprised a total of 10,000 possible sequences in our behavior test. It is unlikely that the monkeys remembered every sequence and its associated value. In other words, their brain most likely did not establish an association between each sequence and its corresponding reward. It is conceivable that with enough training, the monkeys might finally memorize all the sequences, and the OFC presumably would encode the value associated with each sequence as a result.

Such reasoning may be extended to the situation in which an animal tries to assess a concrete object that is unfamiliar to it such as a novel food item. Because there is no established value association yet, the animal has to resort to other means to assess its value, possibly by examining its individual features such as color, shape, and smell. In this case, we would like to argue that the OFC is not directly involved in the calculation of its value initially. Instead, it encodes the value of the individual features that the brain is familiar with. The DLPFC may play the role of integrating information from these features and calculate the object’s value. After the animal gaining enough experience, the OFC may start to encode the value associated with the object as a whole.

### OFC and Sequential Processing

Several recent studies point to the possibility that the OFC processes value information in a sequential manner, which may be guided by attention (McGinty et al., 2016; Rich and Wallis, 2016; Xie et al., 2018). Notably, Rich and Wallis showed that the OFC neural activity alternates between representing the value of each option during decision making. In our study, the stimuli were presented sequentially. Still, our finding of the lack of sustained encoding of the stimulus value or the accumulated evidence in the OFC adds to the body of evidence supporting that the OFC encodes value information in a sequential manner and the integration of information occurs outside the OFC. In this sense, the OFC may be regarded as an extension of the ventral stream of the visual system, which further translates object identity information into its behavior relevance.

### Mixed selectivity

We created a neural network model in order to understand how value information regarding to color is transformed into the action domain in the DLPFC. The results suggested that neurons with mixed selectivity were essential. The importance of such mixed selectivity has been demonstrated both experimentally and theoretically (Barak et al., 2013; Blanchard et al., 2018; Cheng et al., 2015; Rigotti et al., 2013; Zhang et al., 2018). Our results again confirmed the existence of neurons with mixed selectivity in the DLPFC. In addition, we provided clues on the computation underlying their mixed selectivity.

### DLPFC vs. LIP

Previous studies using a similar behavior paradigm showed that the LIP neurons also encoded the combined evidence (Kira et al., 2015; Yang and Shadlen, 2007). The similarities between the DLPFC and the LIP were also observed in experiments using random dot motion discrimination tasks(Kim and Shadlen, 1999; Roitman and Shadlen, 2002). The new results again raise the question of the relative roles that the DLPFC and the LIP play in decision making.

Here we reported that the DLPFC neurons were found to encode single weights in both the color and the action domain in addition to the summed weights in the action domain. On the surface, the DLPFC by itself seems to possess all the necessary pieces for stimulus-to-action computation. If this is true, the decision-making signal found in the LIP may be inherited from the DLPFC. A recent study that failed to find decision making impairments in monkeys with LIP lesions provided additional support to this argument. (Katz et al., 2016).

However, we do not believe that is the case. The choice signal that we observed in the DLPFC started to reach significance 1280 ms after the first shape onset (Figure 3B). This latency was much larger than that found in the LIP, which was reported to be ~150-200 ms after the shape onset in a study that used a very similar behavior paradigm (Yang and Shadlen, 2007). The very long latency of the choice signal in the DLPFC, if verified, would be too late to contribute to decision making. Such long latency was not observed in a comparable previous study (Kim and Shadlen, 1999). The difference between our results and the Kim and Shadlen study may be understood if we interpret the observed choice signal in the DLPFC as a motor related signal that would only start ~1.5 sec before the actions. The trials in our study were much longer than those in Kim and Shadlen. The choice actions occurred at more than 2.5 sec after the shape onset. Thus, the onset of the observed choice signal was also pushed back a lot compared to the random dots task used in Kim and Shadlen.

We admit that the measure of latency is noisy and may succumb to statistical artifacts. Yet, it is undeniable the representation of the decision variable in the LIP is more consistent and less heterogeneous than that in the DLFPC. Given the extensive connections between the DLPFC and the LIP, it is entirely possible that the LIP is where the integration first occurs and the DPLFC inherits the results for the purpose of motor preparation. Future investigations are still required to distinguish the roles that the two areas play in decision making.

### Summary

Our study explored the roles of the OFC and the DLPFC play in decision making. We discovered that the computation of decisions of eye movements is carried out entirely in the action domain. We further provided the evidence supporting the hypothesis that the DLPFC is where the stimulus-to-action transition occurs. We showed that the OFC encoded value in a transient manner and did not integrate information across time. Taken together, our results showed that the OFC and the DLPFC play distinct roles during value-based decision making.

## Methods

### Subjects and Materials

Two naïve male rhesus monkeys (Macaca mulatta) were used in the study (K and E). They weighed on average 6-7 kg during the experiments. All procedures followed the protocol approved by the Animal Care Committee of Shanghai Institutes for Biological Sciences, Chinese Academy of Sciences (Shanghai, China).

In each experimental session, the monkeys were seated in a primate chair viewing a 23.6-inch video monitor, which was placed at 60 cm distance. An infrared oculometer system (EyeLink 1000) was used to monitor the eye positions at a sampling rate of 500 Hz. Juice or water reward was given to the monkeys based on their preference. The liquid delivery was controlled by a computer-controlled solenoid. The monkeys drank ~150-250 ml per experimental session.

### Behavioral Task

We trained two monkeys (K and E) to perform a probabilistic reasoning task. The monkeys started each trial by fixating and maintaining their gaze on a central fixation point (FP) (0.2° in diameter) on a computer monitor. After the monkeys acquired fixation for 500 ms, a green and a red target showed up on the left and right side of the FP at the eccentricity of 6°. Both colors could appear on either side, which was randomly selected from trial to trial. After another 500 ms, four shapes were shown sequentially near the FP. For monkey K, the center of the shapes was the same as the FP, while for monkey E, the center of the shapes was at a random location chosen from the 4 vertices of an invisible 1° by 1° grid centered on the FP. The shapes were white line drawings and approximately 1.5° by 1.5°. Each shape was presented for 300 ms. Between two consecutive shape presentations, there was a 200 ms delay in which only the FP and the targets were on the screen. Thus, each shape epoch was 500 ms long. The FP disappeared 700 ms after the offset of the forth shape, instructing the monkeys to report their choice. The monkeys had to make a saccadic eye movement toward one of the targets within 1 sec and hold their fixation on it for another 560 ms. The juice reward would be delivered at the end of the fixation of the target.

The reward was determined probabilistically. The probabilities of getting a reward by choosing the red and the green target were:

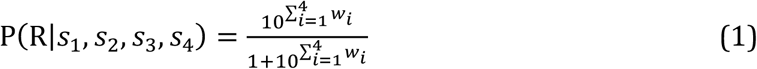

and

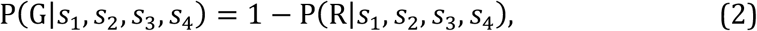

where *s_i_* represents the shape shown in the *i*-th epoch, *w_i_* represents the weight assigned to *s_i_*, and P(R|*s*_1_, *s*_2_, *s*_3_, *s*_4_) and P(G|*s*_1_, *s*_2_, *s*_3_, *s*_4_)are the reward probabilities of the red target and the green target given the shape sequence *s*_1_, *s*_2_, *s*_3_, *s*_4_, respectively. P(R|*s*_1_, *s*_2_, *s*_3_, *s*_4_) and P(G|*s*_1_, *s*_2_, *s*_3_, *s*_4_) add up to 1. The infinitive weights with opposite signs may cancel each other. The reward probability for the red target is 1 with non-cancelled +∞ shape sequences and is 0 with non-cancelled -∞ shape sequences.

### Surgery

The monkeys received a chronic implant of a titanium headpost with standard procedures before the training. After their performance reached a satisfactory level, we performed a second surgery to implant an acrylic recording chamber over the prefrontal region, inside of which a craniotomy was made. The chamber had an inner size of 19.5mm×24mm and was centered over the left principle sulcus. All surgery procedures were done under aseptic conditions. The monkeys were sedated with ketamine hydrochloride (5–15 mg/kg, i.m.) and anesthetized with isoflurane gas (1.5–2%, to effect). Their body temperature, heart rate, blood pressure, and CO2 were monitored during the surgeries.

### MRI

Before and after the recording chamber was implanted, we scanned the monkeys with a Siemens 3T scanner to identify and verify recording locations. The monkeys were sedated with ketamine hydrochloride (5−15 mg/kg, i.m.) and anesthetized with isoflurane gas (1.5−2%, to effect) during the scanning.

### Behavioral Analyses

We computed the percentage of choosing red target and fit a psychometric curve with the least square method:

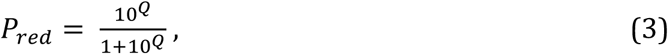

where 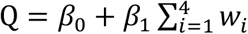. Trials with the shapes of infinite weights were excluded. Unless otherwise mentioned, these trials were also excluded in all the analyses here after.

To test the effects of individual shapes on the monkeys’ choices, we applied a logistic regression, where the regressors were the appearance counts of each shape presented in a trial:

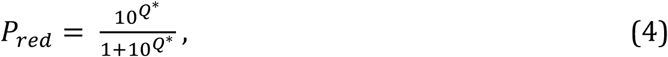

where 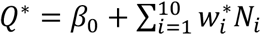, and *N_i_* is the appearance count for the *i*-th shape. We defined the fitted coefficients 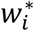 as the subjective weights. All 10 shapes were included in this analysis.

The behavior analyses were based on the same sessions of the electrophysiology recordings used in the analyses below.

### Electrophysiology

The recording procedures were described in our previous study (Xie et al., 2018). Briefly, neuronal responses were recorded using single electrodes (FHC or AlphaOmega) with an AlphaLab SnR System (AlphaOmega). Only units with reasonable isolated wave forms were recorded. Offline sorting was used to further improve the data quality (NeuroExplorer). 2-4 single electrodes were used in each session. The microelectrodes were driven by a multi-channel micromanipulator (Alpha Omega EPS).

We recorded single unit activities from 277 cells in the OFC (121 and 156 from monkeys K and E, respectively), and 384 cells in the DLPFC (170 and 214 from monkeys K and E, respectively). According to MRI results and the neural activities observed during penetrations, the OFC recording locations were on the ventral surface of the frontal lobe between the lateral and medial orbital sulci, roughly corresponding to Walker’s areas 11 and 13 (Walker, 1940). The DLPFC recording were from both banks of the posterior portion of the principal sulcus, in the Brodmann areas 9 and 46d.

### Example neuron PSTH

The firing rate of the example neurons was calculated with a 200ms sliding window. The trials were sorted into 4 quartiles by the variable under discussion (summed spatial weight in Figure 2A and single color weight in Figure 2B) in each epoch.

### Choice Analyses

To see how the neuronal activities in the DLPFC and OFC reflected the monkeys’ choice, we calculated the firing rate differences of each neuron between the trials of different choice outcomes (Figure 3). Specifically, for each neuron, we first grouped the trials by the final choices regarding to either the color or the eye movement direction. The average firing rate for each choice was then calculated. We defined the choice with the larger mean firing rate within the time window from 2 - 2.5 sec after the first shape onset, which was the delay period after the presentation of the last shape and before the offset of the FP, as the neuron’s preferred choice. Then we plotted mean firing rate difference between the preferred and the non-preferred choice across the population. For the control, we shuffled the label of choices and repeat this procedure for 100 times. The significance was tested with a one-way analysis of variance (ANOVA) with unbalanced data at every time point between the experimental data with each unit as a sample and the shuffled data with each unit in each shuffle as a sample. The significance level was 0.01 and no correction of multiple comparison was used. The latency of the choice signal was defined as the first time point when a continuous significant choice signal longer than 100 ms was found.

Because of how we defined the choice preference, even for the shuffled data, the difference between the preferred and the nonpreferred choices was larger than 0.

### Linear regression

We performed linear regressions on four variables for each shape epoch: single color weight (*Wc*), summed color weight (Σ*Wc*), single action weight (*Wa*), summed action weight (Σ*Wa*). The single and summed weights were the same in the first epoch by definition. For each neuron, we convoluted its firing rate (*FR*) with a 200ms square wave at 10 ms steps. We then regressed the *FR* at each time step on the respective variable in epoch *i* (*Factor_i_*).

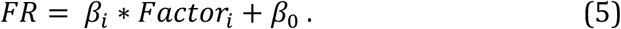

Note that the regression covered the whole trial length when the particular factor was only associated with the shape presented in epoch *i*. The results preceding the epoch *i* reflect the noise level, while the results following the epoch *i* onset reflect how the information was sustained.

### Normalized ∆*R*^2^

The normalized ∆*R*^2^ was calculated similarly to the procedure described by Cai and Padoa-Schioppa (Cai and Padoa-Schioppa, 2014). Briefly, to obtain ∆*R*^2^, we first subtract the *R*^2^ of the regression model in Eq. 5 by the *R*^2^ of the shuffled regression model in which the pairing of the *FR* and *Factor_i_* was shuffled. In order to convert ∆*R*^2^ into a normalized Z score, we shuffled the spike count in a control baseline period (300 – 0 ms before the target onset) across all trials for 1000 times. We calculated the ∆*R*^2^ for the shuffled samples and then obtained the distribution for the baseline 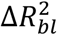. For each neuron and each factor *i*, the normalized ∆*R*^2^ was:

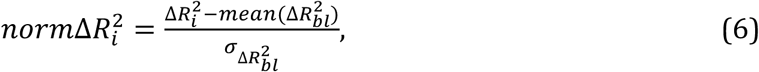

where mean (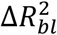) and 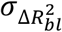 are the mean and the standard deviation of (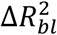), respectively. Following the same procedure, we calculated the normalized population average of ∆*R*^2^ by further normalizing the *norm*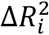:

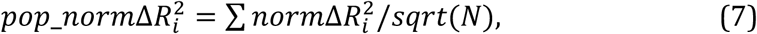

where *N* is the number of neurons. Thus, the population ∆*R*^2^, at the chance level, has an expected value of 0 and a standard deviation of 1, and is comparable across different populations or brain areas.

A one-tailed t-test was used to test the significance of the difference between the population normalized ∆*R*^2^ and 0. In addition, a cluster-size-based thresholding method was used to address the multi-comparison problem (Forman et al., 1995). Specifically, a continuous period is considered as significant if and only if the p-value computed from the t-test at every time point in this period is smaller than 0.1995 and the length of this period exceeds 270ms. The combination of the *p* threshold (*p*<0.1995) and the cluster size threshold (t>270ms) holds the probability of falsely detecting a significant cluster from noise at 0.05.

The latency of a particular variable was defined as the first time point of the first cluster that was found to be significant.

### Lasso

To alleviate the effects of inter-dependency of the variables on the simple linear regression analyses, we created a full model using LASSO (Tibshirani, 1996) that contained the single weight, the summed weight and the choice regressors:

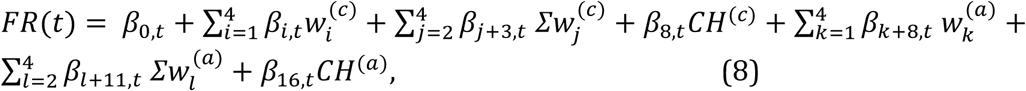

where *FR*(*t*) is the firing rate of the neuron at time t, *β_i,t_* are the fitted coefficients at time 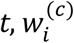 and 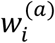 represent the individual weights associated with the shape in the *i*-th epoch in the color and the action domain, respectively, 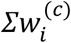 and 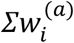 represent the summed weights in the *i*-th epoch in the color and the action domain, respectively. Thus, 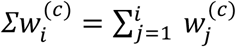 and 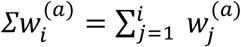. Note that *∑W*_1_ = *W*_1_ therefore we did not include *∑W*_1_ in the model. 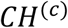 and 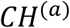 are the choices in regard to the color and the eye movement direction, respectively.

While the summed weights are completely dependent on the single weights, adding an L1-norm penalty of the coefficients in the loss function encourages the fitting algorithm to use a smaller number of regressors, thus the LASSO model biases toward the summed weights (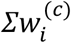 and 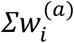) when they can explain the data by themselves.

The LASSO model was fitted independently for each neuron at each time point. All the vectors of regressor samples were normalized into unit vectors before model fitting. We used the built-in function of MATLAB, *lasso*, to fit our data. A 10-fold cross validation procedure is used to determine the regularization parameter *λ*. Specifically, we grid searched *λ* in the log space, from 10^-2^ to 10^0.6^ by a step of 10^0.2^. For each single unit, we found the *λ* that made the most time points to have the smallest mean squared error and used the *λ* as the penalty parameter of this unit.

To estimate the effect size of each factor, we use the absolute value of the standard regression coefficient computed with the following equation:

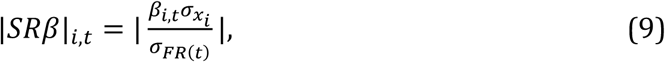

where 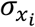 is the standard deviation of the *i*-th regressor, and *σ_FR_*_(*t*)_ is the standard deviation of *FR(t)*. Again, to make the |*SRβ*| comparable across different neuronal populations, we followed the normalization procedure described by Cai and Padoa-Schioppa (Cai and Padoa-Schioppa, 2014). An 800 ms time window before the first shape onset was used as the control baseline period, which was shuffled 100 times to obtain the mean and the standard deviation of the *SRβ*. The significance of the normalized |*SRβ*| was determined similarly as the analysis above for the normalized ∆*R*^2^ in the linear regressions.

### Neural Network Model

The neural network had two input units, one output neuron, and one hidden layer with *tanh* units. The feedforward rule of this network was described by the following equations:

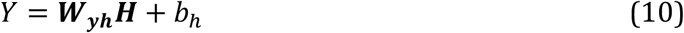

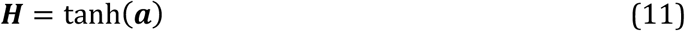

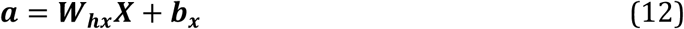

Here, the symbols in bold represent matrices or vectors. Y is the output neuron activity. ***W_yh_*** is a 1-by-*n* connection matrix representing the connection weights from the hidden layer units to output neuron, and *n* is the number of hidden layer units. ***H*** is an n-by-1 vector representing the activities of the hidden units, ***a*** is an n-by-1 vector representing the state of the hidden units, ***W_hx_*** is an n-by-2 connection matrix representing the connection weights from input ***X*** to the hidden layer, and ***X*** is a 2-by-1 vector consists of a single color weight (ranging from −1 to 1) input unit and a target configuration (+1 or −1) input unit. Both *b_h_* and ***b_x_*** are bias terms. A gradient-descent algorithm was used to minimize the mean squared error between the output of the network and the target output (*Wa*).

We started testing with n=16 hidden layer units and gradually reduced the number until 2, with the network performance being maintained. The reduced network helped us to understand how the network worked. By studying the connection weights and the bias terms in the trained model, we extracted its key features depicted as in Figure 8A. The reduced network could be described by the following equations:

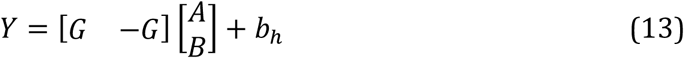

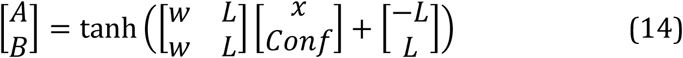

where *Y* is the output of the network (*Wa*), input *x* is the single color weight (*Wc*), *Conf* is the target configuration, *A* and *B* are the activities of the two hidden units, *G* is the connection weight between the hidden units and the output, *b_h_* is a bias term added to the output unit *Y*, *L* is a bias term that is large enough to saturate the *tanh* function, and *w* is the connection weight between the input *x* and the hidden layer units that is small enough so that tanh(*wx*) ≈ *wx*. We set *L*=100, *w*=0.1, *G*=*b_h_*=10.

### Mixed Selectivity Analysis

We used response variance to quantify the mixed selectivity of the DLPFC neurons. The hypothetic neurons are not sensitive to the *Wc* under one spatial configuration, which lead to a small response variance. Their response variance is much larger under the other spatial configuration when they are modulated by the *Wc*. We defined a variance difference index *VDI* as:

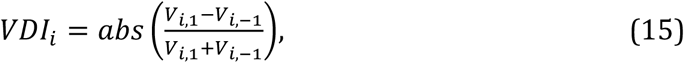

where *V_i_*_,±1_ is the variance of the normalized responses of neuron *i* to the different shape cues when the target configuration equals ±1, respectively. To compute the *V_i_*_,±1_ of a neuron, we followed these procedures. We counted the spikes within an 800 ms time window after each shape’s onset and sorted them by the shape identity and the target configuration. We normalized the responses by subtracting the minimal response and then dividing them by the range:

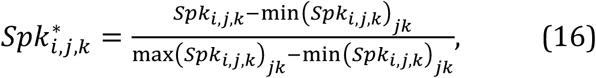

where *Spk_i,j,k_* is the averaged spike count of neuron *i* to shape *j* and under configuration 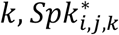 is the normalized response, min(…)_*jk*_ and max(…)_*jk*_ are the minimum and maximum values across all combinations of shapes and configurations.

The variance of response for neuron *i* and configuration *k* is then computed:

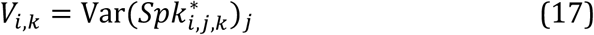

Var(…)_*j*_ is the operation computing the variance across all shapes.

With *V_i,k,_* we computed *VDI_i_* for each neuron *i* based on equation 15. Large *VDI’*s indicate a response pattern similar to that of the hidden units in the network model. Shapes with infinitive weights were included in this analysis.

## Acknowledgements

We thank Yang Xie, Zhewei Zhang, Wenyi Zhang, and Wei Kong for their help in all phases of the study, and Xinying Cai for providing comments and advice. The authors declare no competing financial or nonfinancial interests.

